# JMJD6 Cleaves MePCE to Release P-TEFb

**DOI:** 10.1101/725853

**Authors:** Schuyler Lee, Haolin Liu, Ryan Hill, Xia Hong, Xinjian Liu, Fran Crawford, Qianqian Zhang, Molly Kingsley, Zhongzhou Chen, Andreas Lengeling, Kathrin Bernet, Philippa Marrack, John Kappler, Kirk Hansen, Qiang Zhou, Chuan-Yuan Li, Gongyi Zhang

## Abstract

More than 30% of genes in higher eukaryotes are regulated by promoter-proximal pausing of RNA polymerase II (Pol II). Phosphorylation of Pol II-CTD by positive transcription elongation factor (P-TEFb) is a necessary precursor event that enables productive transcription elongation. The exact mechanism on how the sequestered P-TEFb is released from the 7SK snRNP complex and recruited to Pol II-CTD remains unknown. In this report, we reveal methylphosphate capping enzyme (MePCE), a core component of the 7SK snRNP complex, as the cognate substrate for Jumonji domain-containing 6 (JMJD6)’s novel proteolytic function. Our evidences consist of a crystal structure of JMJD6 bound to methyl-arginine, enzymatic assays of JMJD6 cleaving MePCE *in vivo* and *in vitro*, binding assays, and downstream effects of *Jmjd6* knockout and overexpression on Pol II-CTD phosphorylation. We propose that JMJD6 assists bromodomain containing 4 (BRD4) to recruit P-TEFb to Pol II-CTD by disrupting the 7SK snRNP complex.

## Introduction

Mechanisms of transcription regulation in bacteria are very well established; transcription factors bind to specific DNA to recruit RNA Polymerases (RNAP) to carry out transcription (Ptashne and Gann 1997, Zhang, Campbell et al. 1999). In eukaryotes, however, there are additional layers of regulation such as the nucleosome structures, which could prevent RNA Polymerases including RNA Polymerase I (Pol I), RNA Polymerase II (Pol II), and RNA Polymerase III (Pol III) from productive transcription due to high binding affinity between DNA and histones. Precisely how RNA Polymerases overcome nucleosomal barriers to undergo a productive transcription elongation and how Pol II pausing is regulated remain unanswered (Zhou, Li et al. 2012, Jonkers and Lis 2015, Core and Adelman 2019). In higher eukaryotes, over ~30% genes are regulated by Pol II promoter-proximal pausing (Core, Waterfall et al. 2008, Nechaev, Fargo et al. 2010, Min, Waterfall et al. 2011), which is resultant from nucleosome barriers at +1 position of transcription start sites (Gilchrist, Dos Santos et al. 2010, Weber, Ramachandran et al. 2014, Voong, Xi et al. 2016). We recently discovered that a group of JmjC domain containing protein family including JMJD5 and JMJD7 specifically cleave histone tails and potentially generate tailless nucleosomes. The cleavage activity by JMJD5 and JMJD7 could be associated with the release of the promoter-proximal paused Pol II and trigger Pol II into productive elongation in higher eukaryotes, such as mouse and human (Liu, Wang et al. 2017, Liu, Wang et al. 2018). The cleavage activity of JMJD5 on histone tails was also independently reported by another group (Shen, Xiang et al. 2017), thus cross-validating our respective discoveries.

Compared to efficient recruitment of P-TEFb (including CDK9 and Cyclin T1) by TAT protein in human immunodeficiency virus (HIV)(Peterlin, Brogie et al. 2012), BRD4 is claimed to be responsible for the recruitment P-TEFb to the promoters of Pol II pausing regulated genes (Jang, Mochizuki et al. 2005, Yang, Yik et al. 2005, AJ, Bugai et al. 2016). However, the binding affinity between BRD4 and P-TEFb (~0.5μM) (Itzen, Greifenberg et al. 2014) is much weaker than that that of TAT and P-TEFb (~3nM) (Wei, Garber et al. 1998, Tahirov, Babayeva et al. 2010, Schulze-Gahmen, Lu et al. 2014), and BRD4 is lacking a RNA binding motif (Wu et al. 2007). Therefore, we hypothesize there must exist another factor to help BRD4 to recruit P-TEFb and engages in the instigation of Pol II transcription elongation. Besides the classic Bromo-domains which recognize acetylated histone tails, BRD4 contains an extra terminal domain (ET) recognizing JMJD6 (Rahman, Sowa et al. 2011, Konuma, Yu et al. 2017). Incidentally, we found that JMJD6 nonspecifically binds to single stranded RNA with high affinity (~40nM)(Hong, Zang et al. 2010). We propose that JMJD6 may be recruited by both BRD4 and newly transcribed RNAs from Pol II to help BRD4 recruit P-TEFb, acting analogously to that of TAT protein associating with both P-TEFb and TAR.

JMJD6 is one of the most controversial proteins in biology (Vangimalla, Ganesan et al. 2017). It was first cloned as phosphatidylserine (PS) receptor (Fadok, Bratton et al. 2000), but was corrected as a nucleus expressed protein unrelated to PS (Bose, Gruber et al. 2004, Cikala, Alexandrova et al. 2004, Cui, Qin et al. 2004). It was later reported to contain arginine demethylase activity on histone tails (Chang, Chen et al. 2007), hydroxylase activity on splicing factor U2AF65 (Webby, Wolf et al. 2009) and histone tails (Han, Li et al. 2012), and both arginine demethylase activities on histone tails and RNA demethylase activities on 5’ prime of 7SK snRNA (Liu, Ma et al. 2013), and surprisingly PS binding (Neumann, Coakley et al. 2015, Yang, Chen et al. 2015). The exact or cognate substrate(s) of JMJD6 remains unresolved or controversial. Based on the novel protease activities of JMJD5 and JMJD7 (Liu, Wang et al. 2017, Shen, Xiang et al. 2017, Liu, Wang et al. 2018), the high structural similarity among catalytic cores of JMJD5, JMJD6, and JMJD7 (Hong, Zang et al. 2010, Liu, Wang et al. 2018), and analogous severe phenotypes among knockouts of *Jmjd5* and *Jmjd6* in mice (Li, Sarkisian et al. 2003, Bose, Gruber et al. 2004, Ishimura, Minehata et al. 2012, Oh and Janknecht 2012), we hypothesized that JMJD6 may contain protease activity working on methylated arginines on some protein candidates which regulate the activity of Pol II, especially promoter-proximally paused Pol II. In this report, we reveal that methylphosphate capping enzyme (MePCE) (Jeronimo, Forget et al. 2007, Xue, Yang et al. 2010) of the 7SK snRNP complex is a cognate substrate of JMJD6.

### JMJD6 has a unique structure to hold methyl-arginine

Based on these divergent reports regarding substrates of JMJD6 (Chang, Chen et al. 2007, Webby, Wolf et al. 2009, Han, Li et al. 2012, Liu, Ma et al. 2013, Neumann, Coakley et al. 2015), we re-interrogated proposed substrates using stringent and unified criteria. As we reported previously, JMJD6 binds with high binding affinity (~40nM) to single stranded RNA (ssRNA) without sequence specificity (Hong, Zang et al. 2010). However, truncation analysis showed that JMJD6 barely binds to ssRNA without the C-terminal flexible region (Hong, Zang et al. 2010). This suggests that the C-terminal domain of JMJD6 may just serve as ssRNA binding motif and RNAs are not a substrate for the enzymatic activity of JMJD6. On the other hand, the structure of the catalytic core of JMJD6 shows some critical similarity to those of JMJD5 and JMJD7, with a negatively charged microenvironment near the catalytic center (Hong, Zang et al. 2010, Liu, Wang et al. 2018), suggesting positively charged substrates (Fig. 1). As we reported, JMJD5 and JMJD7 specifically recognize methylarginines of histone tails *via* a Tudor-domain-like structure near the catalytic center of JMJD5, which could specifically recognize methylarginines, but not methyllysine (Liu, Wang et al. 2017, Liu, Wang et al. 2018). We reasoned that the similar structural features among JMJD6, JMJD5, and JMJD7 may confer similar substrate for JMJD6 as those of JMJD5 and JMJD7. In this regard, crystals of JMJD6 without C-terminal motif (1-343) were soaked with a monomethylarginine derivative. Interestingly, four out of eight JMJD6 molecules within an asymmetric unit bound to monomethylarginine, which coordinates with Fe2+ and alpha-KG in the catalytic center similar to that of JMJD5 and methylarginines (Fig. 1, Fig.S1, Fig. S2, Fig. S3, Table S1). However, the methylated sidechain of arginine is located in a more open catalytic space containing negatively charged residues, compared to that of JMJD5, indicating that the pocket could hold more than one sidechain (Fig. 1A). This may suggest a novel substrate recognition mode, which is different from that of JMJD5 (Liu, Wang et al. 2018). Nevertheless, the complex structure shows several key evidences. First, JMJD6 does bind to substrates with methylarginine or possibly methyllysine or both arginine and lysine with and without methylation (Fig. 1B, 1C). Second, the methyl group is far away from either the divalent ion or alpha-KG, suggesting JMJD6 may not act as lysine or arginine demethylases to remove methyl groups on the sidechain of either lysine or arginine. Third, peptides or proteins could be cognate substrates instead of RNA or DNA, which do not contain any positive charge with or without methylation. Fourth, the catalytic center contains analogous residues present in JMJD5 and JMJD7, suggesting a similar novel catalytic mechanism as those of JMJD5 and JMJD7 through an imidic acid as proton mediator (Fig. 1D)(Lee, Chen et al. 2017, Liu, Wang et al. 2018). This novelty is particularly exemplified by the fact that commercially available protease inhibitor cocktail (Roche) at 2x concentration cannot inhibit activities of JMJD5, JMJD6, and JMJD7. Furthermore, a comprehensive protein composition analysis using mass spectrometry (Table S2) of all purified recombinant JMJD6 samples used in the following experiments could not detect any protease contaminants.

**Figure 1.**
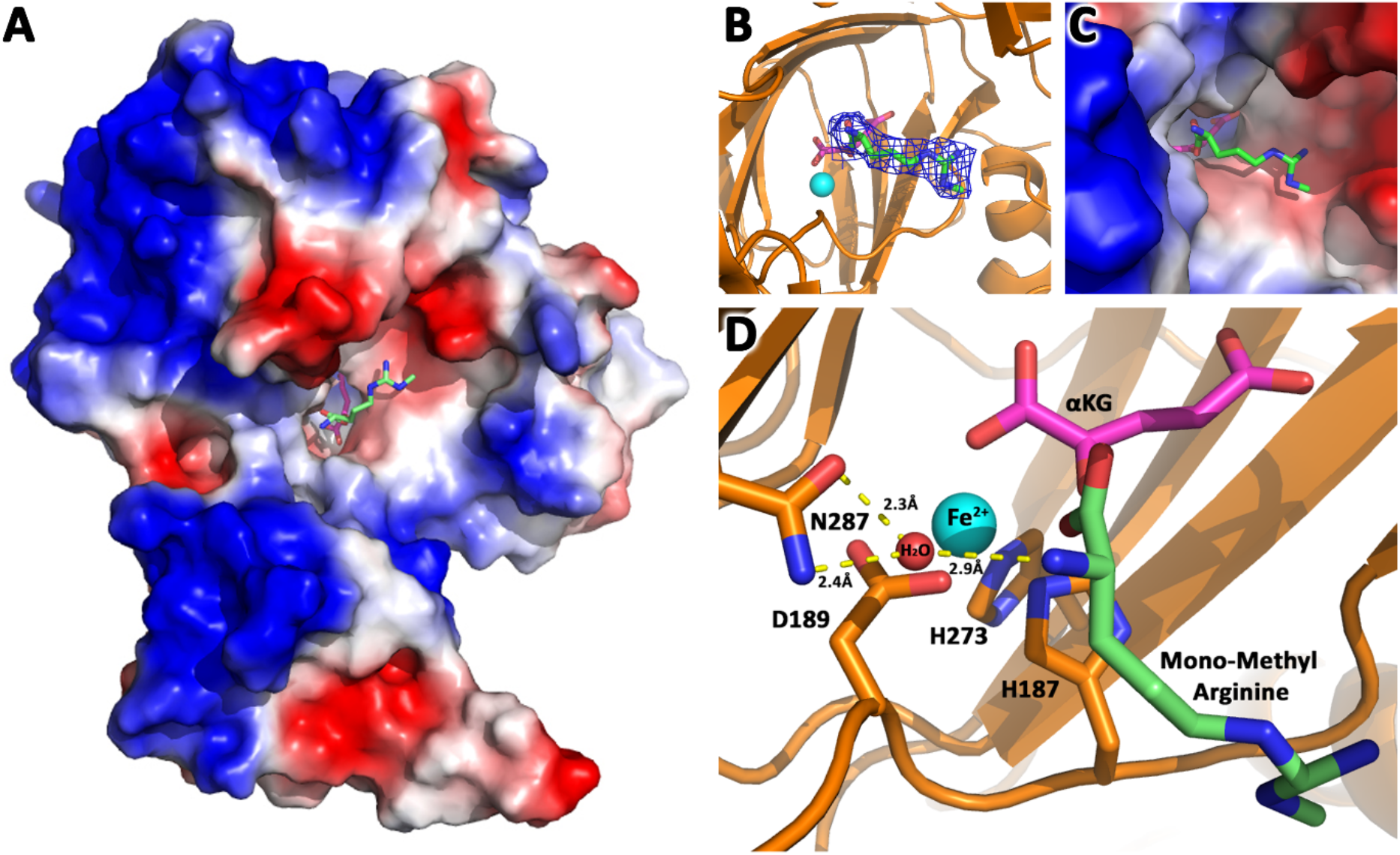
JMJD6 binds to monomethyl arginine (1-of-4). **a.** complex structure of JMJD6 (1-343) and monomethyl arginine (MM-Arg). Surface charges were generated using PyMOL (Action > generate > vacuum electrostatics > protein contact potential) (https://pymol.org/2/). Red represents negatively-charged surface, Gray represents neutral-charged surface, and Blue represents positively-charged surface. **b.** Omit map 2Fo-Fc electron density of MM-Arg. **c.** Magnified view of MM-Arg in the catalytic center of JMJD6 **d.** Coordination of elements at catalytic center.

### JMJD6 cleaves MePCE

Since JMJD5 and JMJD7 make cleavage on histone tails (Liu, Wang et al. 2017), we asked whether JMJD6 also recognizes histone tails. Interestingly, JMJD6 does in fact have activities on bulk histone *in vitro* (Fig. S4A). This result may explain why two groups found that JMJD6 could reduce methylarginine containing histone tails in *in vitro* assays probed with antibodies (Chang, Chen et al. 2007, Liu, Ma et al. 2013), in which post-cleft short peptides with methylarginines can not be recognized by antibodies in western blots. To confirm or rule out whether histone tails are cognate substrates of JMJD6, we first assessed the binding affinity between JMJD6 and histone peptides with or without arginine methylation. From fluorescence polarization binding assays, binding affinities between JMJD6 and peptides are weak and around ~150μM (Fig. S5D, S5E). Most importantly, with or without *Jmjd6*, the level of arginine methylated histones or overall histone levels does not change in MEF cells *in vivo* (Fig. S4B), which is in stark contrast to those of JMJD5 and JMJD7 (Liu, Wang et al. 2017). These data suggest that histone tails are not cognate substrates of JMJD6. Parenthetically, it is reported that JMJD6 binds to LARP7 on 7SK snRNP (Weimann, Grossmann et al. 2013), works on methyl cap of 5’ 7SK snRNA (Liu, Ma et al. 2013), and binds to BRD4 (Rahman, Sowa et al. 2011, Konuma, Yu et al. 2017), all suggesting a close relation with the 7SK snRNP complex.

We hypothesized that JMJD6 may work on some protein component(s) of the 7SK snRNP to regulate the stability of 7SK snRNP complex. Interestingly, all protein components of the 7SK snRNP complex including MEPCE, LARP7, and HEXIM1 are drastically decreased in the MEF cells without *Jmjd6* (Fig. 2A). One possibility is that all members are transcriptionally regulated by JMJD6. This could be consistent with the severe phenotype of *Jmjd6* knockouts, which results in embryonic lethality and cell growth retardation and tissue differentiation (Li, Sarkisian et al. 2003, Bose, Gruber et al. 2004), suggesting that JMJD6 is a global master transcriptional regulator controlling expression of a large group of genes including components from the 7SK snRNP complex. However, our RNA-Seq data do not support direct transcriptional downregulation of 7SK snRNP complex members at mRNA level (Table S3). All of them, including MePCE, LARP7, and HEXIM1 have similar mRNA levels with or without *Jmjd6*. At the moment, the cause of this downregulation of proteins level of these components from 7SK snRNP remains unresolved and is beyond the scope of this report. However, when we introduce back JMJD6 via overexpression into MEF cells lacking *Jmjd6*, protein levels of LARP7 and HEXIM1 are rescued, whereas MePCE is not (Fig. 2A). To account for this disappearance of MePCE, we hypothesized that overexpression of JMJD6 may directly target MEPCE for degradation. To confirm this hypothesis, we respectively overexpressed full-length MePCE in wild-type MEF, *Jmjd6* knockout MEF, wild-type JMJD6 overexpression in *Jmjd6* knockout MEF, and inactive mutant JMJD6 overexpression in *Jmjd6* knockout MEF. The whole cell lysates were probed with anti-MePCE antibody. Our hypothesis was vindicated with the emergence of a lower molecular weight form of MePCE in wild-type MEF and wild-type JMJD6 overexpression in *Jmjd6* knockout MEF, whereas the lower molecular weight form of MePCE was not detectable in the *Jmjd6* knockout MEF, and inactive mutant JMJD6 overexpression in *Jmjd6* knockout MEF (Fig. 2B).

**Figure 2.**
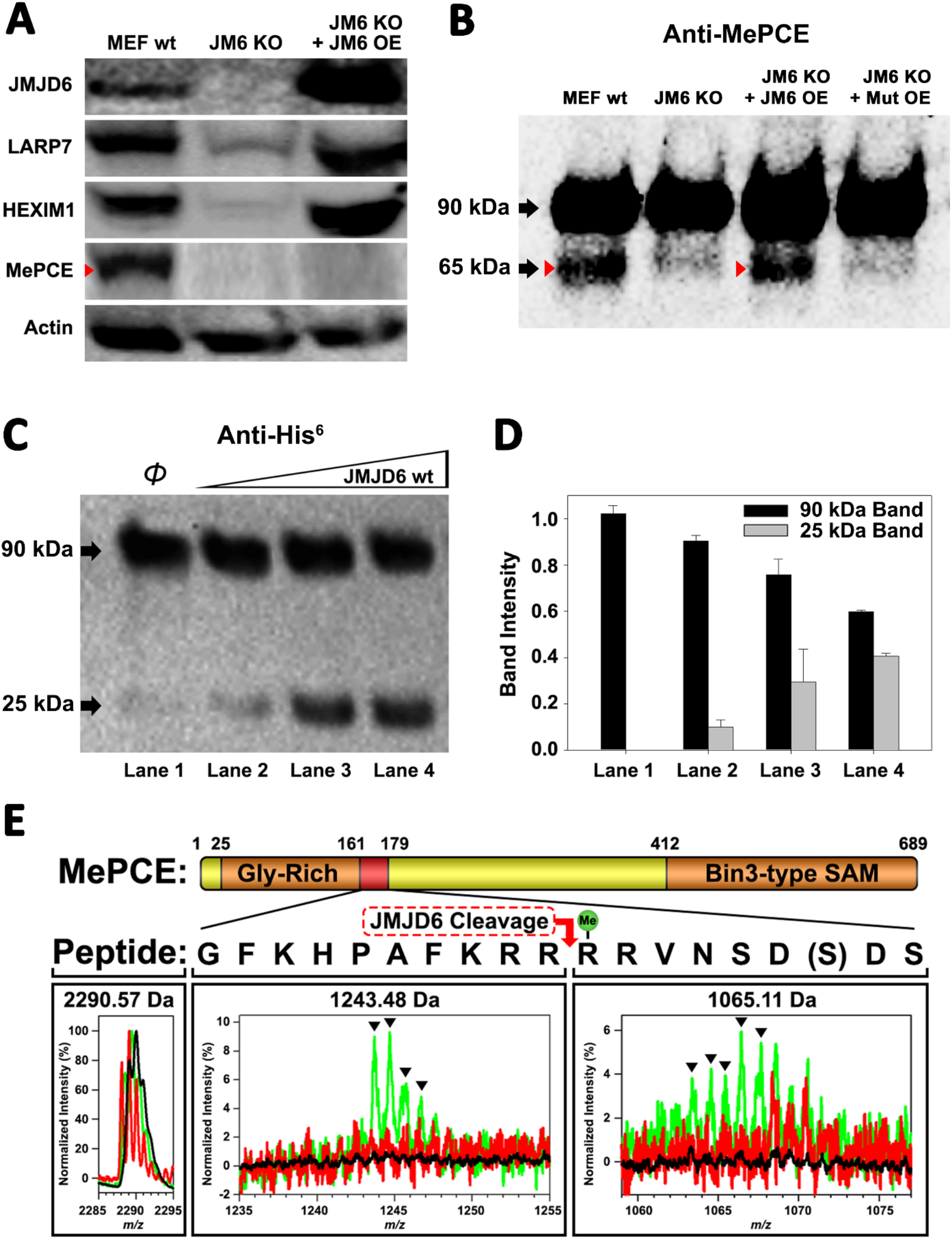
JMJD6 targets MePCE for proteolysis. **a.** Western blot of wild-type MEF, *Jmjd6* knockout MEF, and JMJD6 overexpression in *Jmjd6* knockout MEF probed with antibodies specific for components of the 7SK snRNP complex; LARP7, HEXIM1, and MePCE. **b.** Western blot of MePCE overexpressed respectively in wild-type MEF, *Jmjd6* knockout MEF, wild-type JMJD6 overexpression in *Jmjd6* knockout MEF, and inactive mutant JMJD6 overexpression in *Jmjd6* knockout MEF; probed with antibody specific for MePCE. **c.** Wild-type JMJD6 titrated into full-length MePCE with N-terminal His^6^-tag. Enzymatic activity of JMJD6 is probed with anti-His^6^ antibody. **d.** Quantification of c. **e.** The endopeptidase activity of JMJD6 on synthesized MePCE (161-179) R171-me2s/C177S peptide. The mass spectrum is normalized to the intensity of the undigested peptide input. The peptide is assayed with wild-type JMJD6 (green), inactive mutant JMJD6 (red), or peptide alone (black). The MePCE (161-179) peptide with symmetric dimethylation on R171 and C177S mutation has a molecular weight of 2,290.57 Da. After wild-type JMJD6 cleavage between R170 and R171, the major peaks* (black triangles) with the molecular weight of 1,243.48 Da corresponds to the N-terminal product and the molecular weight of 1,065.11 Da corresponds to the C-terminal product respectively. *The multiple peaks are isotopic distributions, which are characteristic of MALDI-TOF.

To reproduce these *in vivo* results in an *in vitro* setting, full-length proteins linked with N-terminal His^6^-tag of MePCE and HEXIM1 were generated from insect cells and LARP7 and HEXIM2 were generated from bacteria and purified, followed by subjecting these proteins to direct *in vitro* enzymatic assays. Consistent with our *in vivo* results, JMJD6 cleaves MePCE *in vitro* (Fig. 2C. 2D), but not HEXIM1, HEXIM2, nor LARP7 (data not shown). Upon cleavage of MePCE by JMJD6 *in vitro*, a band with molecular weight ~25 kDa was detected by antibody against His^6^-tag at the N-terminal (Fig. 2C). To provide context, MePCE contains a total of 689 residues and a theoretical molecular weight of 74.4 kDa, whereas it is detected at ~90 kDa on SDS-PAGE gel due to its high content of proline residues (10.7%). We expect that the ~25 kDa fragment from N-terminal of MePCE may contain 200 residues or less due to rich proline residues within the N-terminal of MePCE (first 150 residues of MePCE contains 27 prolines, 18.0% prolines). Furthermore, our *in vivo* assay yielded a cleaved MePCE band with a molecular weight ~65 kDa (Fig. 2B). The anti-MePCE antibody used in this assay was generated using a peptide immunogen containing residues 200-250 of MePCE. Thus, the ~65 kDa band may correspond to the cleaved C-terminal segment of MePCE containing the approximate residues 200-689.

Based on the binding of JMJD6 to methylated arginine we obtained from the complex crystal structure, we reasoned that arginine residues within approximately the first 200 residues of MePCE could contain the recognition site. Several peptide fragments including residues from 81 to 160, residues 154 to 184, and residues 187 to 244 were synthesized. Peptides of 81-160 and 187-244 did not show any cleavage when incubated with JMJD6. Peptides of 154-184 showed cleavage activity, but at levels of <1% compared to peptide input. We attributed this low activity as a matter of insolubility and dimerization via C177 oxidation. To overcome this obstacle, a shorter peptide from residue 161 to 179 containing C177S was synthesized and subjected to enzymatic reaction under JMJD6. This peptide also contains symmetric dimethylation on R171, given that our binding data, described in the next section, suggests that this particular modification yields the highest binding affinity (Fig. 3C). In line with our expectations, dominant peaks corresponding to the cleaved MePCE peptide products were detected by mass spectrometry, but not in either peptide alone or with an inactive mutant version of JMJD6 (Fig. 2E).

**Figure 3.**
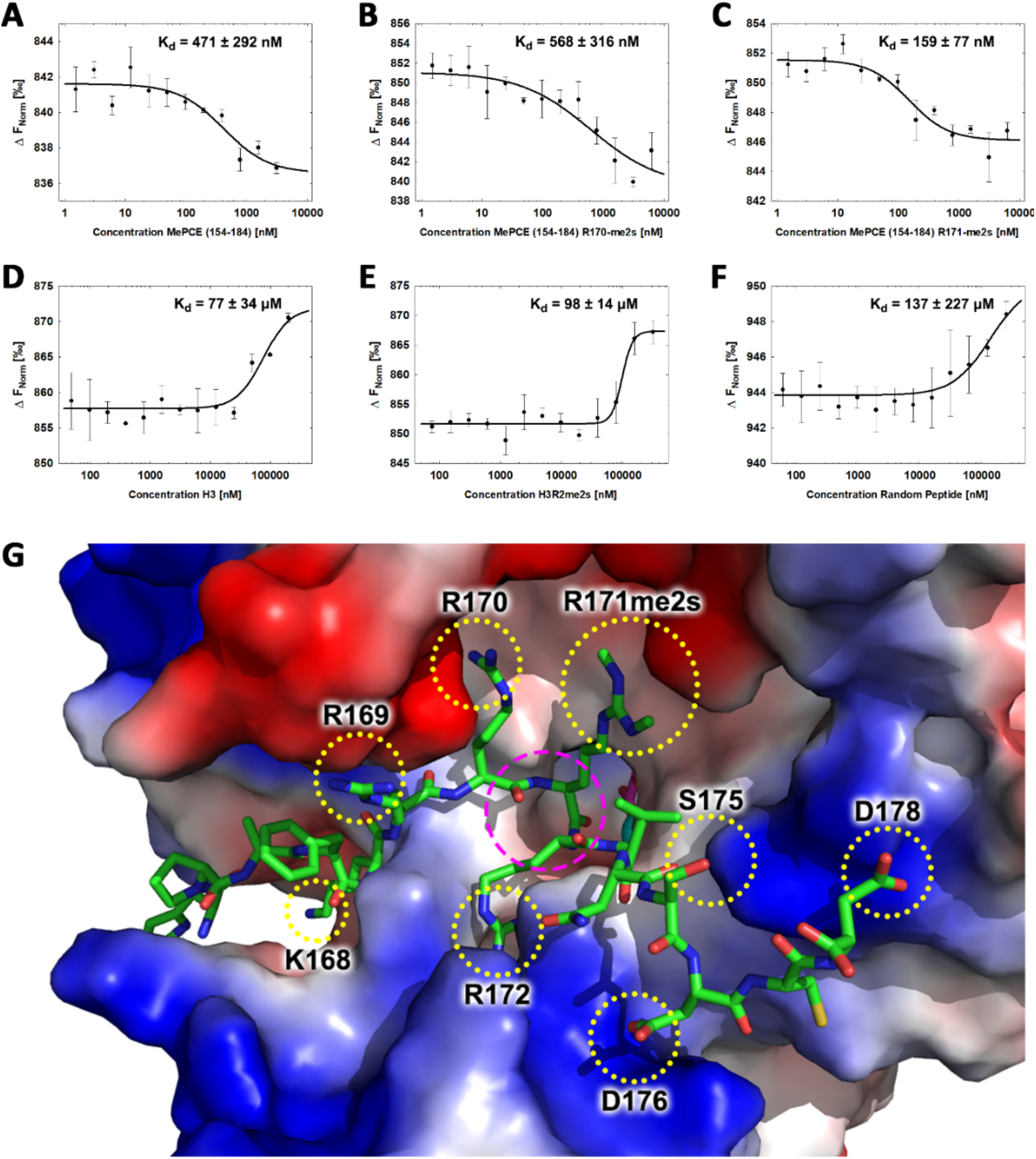
JMJD6 specifically binds to MePCE site containing residues 164-178 (determined via MST). **a**. The binding of His-JMJD6 (1-343) to unmodified MePCE (154-184). **b**. The binding of His-JMJD6 (1-343) to MePCE (154-184) R170-me2s. **c**. The binding of His-JMJD6 (1-343) to MePCE (154-184) R171-me2s. **d**. The binding of His-JMJD6 (1-343) to unmodified Histone 3 (1-21). **e**. The binding of His-JMJD6 (1-343) to Histone 3 (1-16) R2-me2s. **f**. The binding of His-JMJD6 (1-343) to C-peptide (57-87). **g.** Electrostatic interactions between JMJD6 (1-343) and MePCE (164-178) determined by YASARA Energy Minimization server are highlighted in yellow dotted circles and catalytic center is highlighted in magenta dashed circle.

### JMJD6 specifically recognizes methylarginine within MEPCE

To determine the binding affinity between JMJD6 and the newly discerned MePCE proteolysis site, Microscale Thermophoresis (MST) assay was performed using catalytic core of JMJD6 (1-343) titrated with the MePCE (154-184) peptide exhibiting: no modification (Fig. 3A), R170-me2s (Fig. 3B), or R171-me2s (Fig. 3C). The highest binding affinity was exhibited by the MePCE (154-184)-R171-me2s peptide with Kd = 159 ± 77 nM, followed by the peptide containing no modification with Kd = 471 ± 292 nM, and lastly the peptide with R170-me2s with Kd = 568 ± 316 nM. Fluorescence polarization assays were used to cross-validate the Kd values obtained via MST above, all of which respectively fell within the margin of error (Fig. S5A, A5B, S5C). These data suggest that methylation on R171 enhances binding nearly 3-fold compared to no modification, whereas methylation on R170 diminishes binding marginally. Whether or not these particular methylations occur *in vivo* is yet to be explored. Interestingly, the YASARA Energy Minimization Server developed by Dr. Kevin Karplus’ group yielded a theoretical model that positions R171 in the identical position to the monomethyl-arginine observed in our crystal model (Fig. S6A)(Krieger, Joo et al. 2009). Furthermore, the energy-minimized model displayed 8 separate charge-charge interactions between JMJD6 and MePCE (Fig. 3G). Although a *bona fide* complex structure is preferred, this computational model is in excellent agreement with all our findings and provides a coherent justification for the experimentally observed high binding affinities between JMJD6 and MePCE. To compare the abovementioned binding affinities to that of JMJD6 and histone tails, as other groups have previously purported, the identical MST assay was performed using peptides derived from Histone 3 (1-21) containing no modification (H3) (Fig. 3D), Histone 3 (1-16)-R2-me2s (H3R2me2s) (Fig. 3E), and C-peptide (57-87) as a negative control (Fig. 3F). For the Histone 3 peptides, low binding affinity was exhibited with Kd = 77 ± 34 μM for H3 peptide and Kd = 98 ± 14 μM for H3R2me2s peptide. No meaningful binding was observed for the C-peptide (Fig. 3F).

### Knockout of *Jmjd6* leads to down-regulation of Ser2p-CTD of Pol II

We previously reported that cleavage of histone tails by JMJD5 and JMJD7 could lead to tailless nucleosomes, which are unstable and are more readily overcome by Pol II during transcription (Liu, Wang et al. 2017, Liu, Wang et al. 2018). What happens when MePCE is cleaved? Does MePCE cleavage mediate P-TEFb to be released from the 7SK snRNP complex, allowing to be recruited to CTD of Pol II, and then to phosphorylate Ser2-CTD of Pol II? Nuclear extracts from both wild type MEFs, *Jmjd6* knockout MEFs and JMJD6 overexpression in *Jmjd6*-deficient MEFs were subjected to investigation as to the level of Ser2p-CTD of Pol II. Consistent with all our current and previous findings, Ser2p-CTD of Pol II is drastically reduced in samples from MEF cells without *Jmjd6*, despite CDK9 levels remaining unchanged, suggesting that Ser2p-CTD of Pol II is regulated by JMJD6 (Fig. 4A). This result is consistent with the critical role of JMJD6 in embryonic development, which is lethal without *Jmjd6* (Li, Sarkisian et al. 2003, Bose, Gruber et al. 2004). It explains why growth of MEF cells without *Jmjd6* is severely compromised, similar to those without either *Jmjd5* or *Jmjd7* (Liu, Wang et al. 2017). This data suggests not only that JMJD6’s cleavage of MePCE is essential for P-TEFb to be released from the 7SK snRNP complex and ultimately the phosphorylation of Ser2-CTD of Pol II, but also further cross-confirms that a species of Ser2p-CTD of Pol II is indeed generated by CDK9. The content of Ser2p-CTD of Pol II is recovered when JMJD6 is re-expressed (Fig. 4A), suggesting the direct relation of JMJD6 and phosphorylation of Ser2-CTD of Pol II *in vivo.* When these results are aggregated with previously published reports, whereby (1) JMJD6 exhibits strong binding affinity to nonspecific ssRNA as we previously reported (Hong, Zang et al. 2010), (2) JMJD6 associates with BRD4 as reported by others (Rahman, Sowa et al. 2011, Konuma, Yu et al. 2017), and (3) JMJD6 digests MePCE so as to disrupt the overall stability of 7SK snRNP complex as highlighted in this report, a model of P-TEFb transcription regulation unique to higher eukaryotes can be generated: JMJD6 binds to 20-50 nt newly transcribed 5’ prime end of ssRNAs of initiated Pol II of any stimulating genes. The association between BRD4 and JMJD6 allows BRD4 to bring P-TEFb (CDK9) to close proximity of CTD of Pol II, allowing for Ser2-CTD phosphorylation (Fig. 4B). Remarkably, this recruitment of P-TEFb by JMJD6, with the assistance of BRD4, is similar to that of TAT protein hijacking P-TEFb complex after associating with TAR transcript to trigger the expression of HIV retroviral genome (Fig. 4C) (Peterlin, Brogie et al. 2012, Zhou, Li et al. 2012, AJ, Bugai et al. 2016). Here, our observation regarding the direct relation of JMJD6 and level of Ser2p-CTD of Pol II *in vivo* is corroborated by Liu et al.’s report, in which 7SK snRNP complex pulled-down through HEXIM1 antibody from HEK293 cells could be disrupted by JMJD6 *in vitro*, as indicated by the decrease in CDK9 levels (Liu, Ma et al. 2013), thus strongly supporting our proposed role of JMJD6 as a direct disruptor of the 7SK snRNP complex.

**Figure 4.**
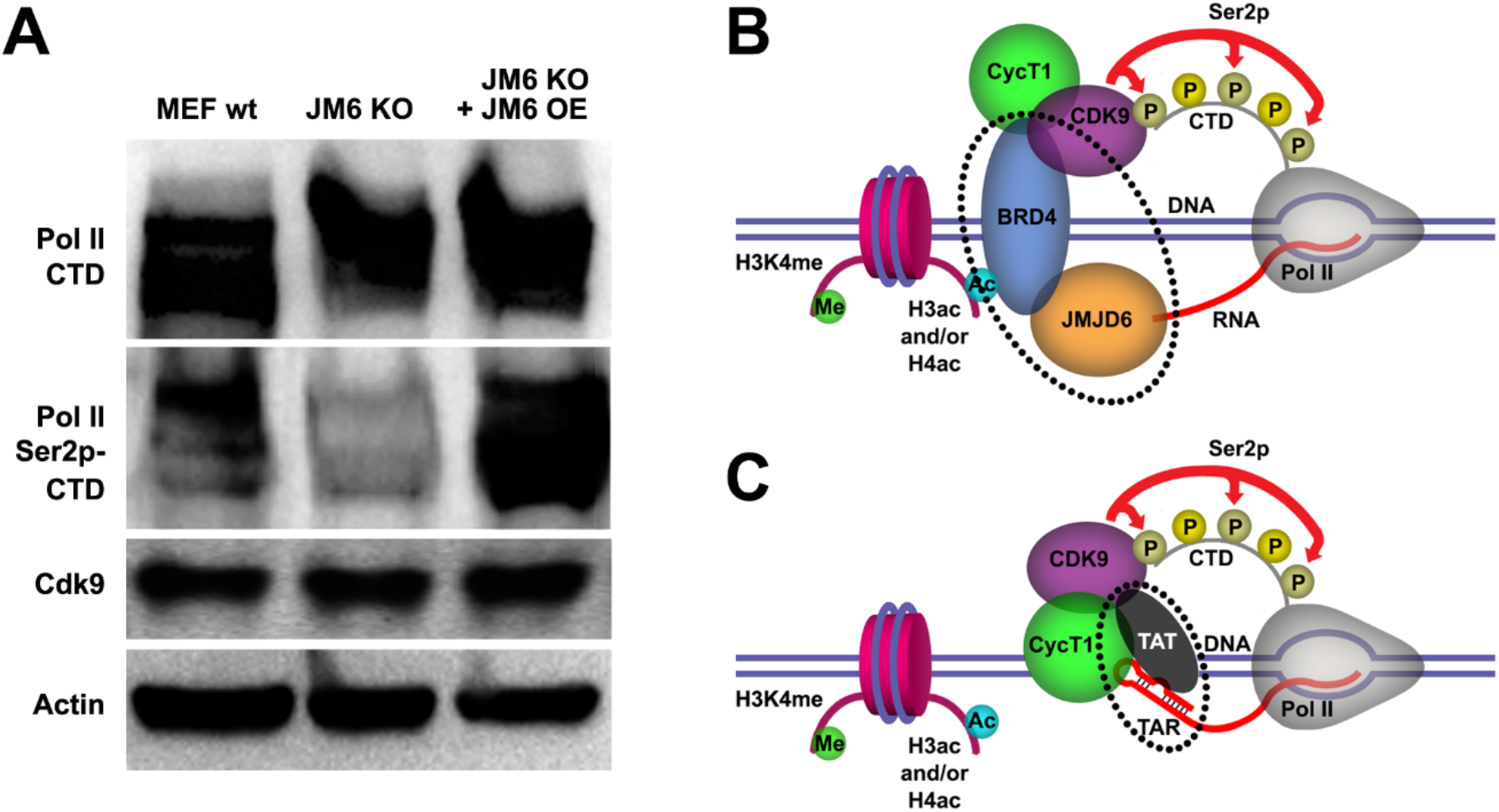
JMJD6 regulates Pol II Ser2-CTD phosphorylation. **a.** Western blot of wildtype MEF, *Jmjd6* knockout MEF, and JMJD6 overexpression in *Jmjd6* knockout MEF probed with antibodies specific for Pol II CTD, Pol II ser2p-CTD, and CDK9. **b.** ssRNA-bound JMJD6 and acetylated H3/H4-bound BRD4 in conjunction (black dotted circle) bridges P-TEFb to paused Pol II. **c.** TAR-bound TAT (black dotted circle) bridges P-TEFb to paused Pol II.

## Discussion

It took us nearly two decades to address the structure and function of JMJD6 since it was first cloned (Fadok, Bratton et al. 2000). During the long journey, based on the conserved structure of JmjC domain containing family, we pioneered in characterizing the catalytic core of JMJD2 subfamily, which turned out to be a novel lysine demethylase (Chen, Zang et al. 2006, Whetstine, Nottke et al. 2006, Chen, Zang et al. 2007). We then solved the structure of JMJD6 with the help of a monoclonal antibody and discovered its unique ssRNA binding property almost a decade ago (Hong, Zang et al. 2010). However, the exact function and its cognate substrate remained a mystery, though several groups claimed variety of enzymatic activities and substrates in past two decades (Chang, Chen et al. 2007, Webby, Wolf et al. 2009, Han, Li et al. 2012, Liu, Ma et al. 2013). The accidental discoveries of novel protease activities of JMJD5 and JMJD7 by our group prompted the exploration of potential protease activity of JMJD6 (Liu, Wang et al. 2017, Liu, Wang et al. 2018). The report by Liu et al. helped us to narrow down the potential cognate substrate within the 7SK snRNP complex (Liu, Ma et al. 2013). Reports of arginine demethylase activity (Chang, Chen et al. 2007, Liu, Ma et al. 2013) helped us to focus on methylated arginine as the potential cleavage sites. Thanks to these reports, we identified MePCE to be the cognate substrate of JMJD6, as well as the potential cleavage site. Numerous lines of evidence from our current discoveries and other publications corroborates the authenticity of this conclusion: First, JMJD6 is specifically associated with BRD4 through the ET domain (Rahman, Sowa et al. 2011, Konuma, Yu et al. 2017), while BRD4 is very well established to recruit P-TEFb to paused Pol II (Yang, Yik et al. 2005, Yang, He et al. 2008). Second, JMJD6 could disrupt the 7SK snRNP complex *in vitro* as reported by Liu et al. (Liu, Ma et al. 2013). Third, our current discoveries showed that JMJD6 cleaves MePCE of the 7SK snRNP complex. Fourth, our previous discovery on the non-specific ssRNA recognition of JMJD6 bridges JMJD6 to the initiated Pol II which generates 20-50 nt long ssRNA (Hong, Zang et al. 2010).

The findings that JMJD5 and JMJD7 are in fact proteases was highly unexpected, and required multiple lines of evidence pursued with great scrutiny. It is quite extraordinary that a very conserved Jumonji domain containing hydroxylase family should contain a subfamily which evolutionarily adopted distinctive protease activities, as well as exhibit several unexpected novelties including the catalysis mechanism involving imidic acid and both endopeptidase and exopeptidase activities (Lee, Chen et al. 2017, Liu, Wang et al. 2017, Liu, Wang et al. 2018). Indeed, recombinant proteins derived from bacteria have a high chance of contamination by bacterial proteases, although we exhausted innumerable means to exclude the possibility of contamination in our assays (Liu, Wang et al. 2017, Liu, Wang et al. 2018). In this report, the novel protease activity of JMJD6, which cleaves *before* the methylated arginine, is remarkably distinct to the cleavage *after* the methylated arginine mediated by JMJD5 and JMJD7 (Liu, Wang et al. 2017, Liu, Wang et al. 2018). This important piece of evidence supports the claim of JMJD5 and JMJD7 as proteases. It is extremely unlikely that one batch of proteins we purified was contaminated by one type of undiscovered protease(s) (cleavage after methylated arginine for JMJD5 and JMJD7) while another batch of proteins was contaminated by a different type of undiscovered protease(s) (cleavage before methylated arginine for JMJD6), all which respectively have substrate specificity matching the expected biological profiles of JMJD5/6/7.

Based on our current discoveries, we may derive a novel transcription regulation pathway for genes regulated by promoter-proximate pausing Pol II. First, upon stimulation of cells, signals will reach specific transcription factors through signal transduction pathways. With modification or with the help of other partner molecules, these transcription factors bind to enhancers close to the paused Pol II. These transcription factors will recruit P300/CBP with the help of H3K4(me1) at enhancer regions, which in turn acetylates H3 and/or H4 on the same nucleosome bound by P300/CBP through association with H3K4(me1) to generate acetylated H3 and/or H4 (Fig. 5A). Next, BRD4 is recruited to acetylated H3 and/or H4, and in turn engages with JMJD6 and the initiated Pol II complex. JMJD6 specifically cleaves MePCE of the 7SK snRNP complex so as to release the P-TEFb complex containing CDK9 (Fig. 5B). Finally, JMJD6 and BRD4 brings P-TEFb to close proximity of CTD of Pol II. P-TEFb (CDK9) then phosphorylates Ser2-CTD of Pol II (Fig. 5C).

**Figure 5.**
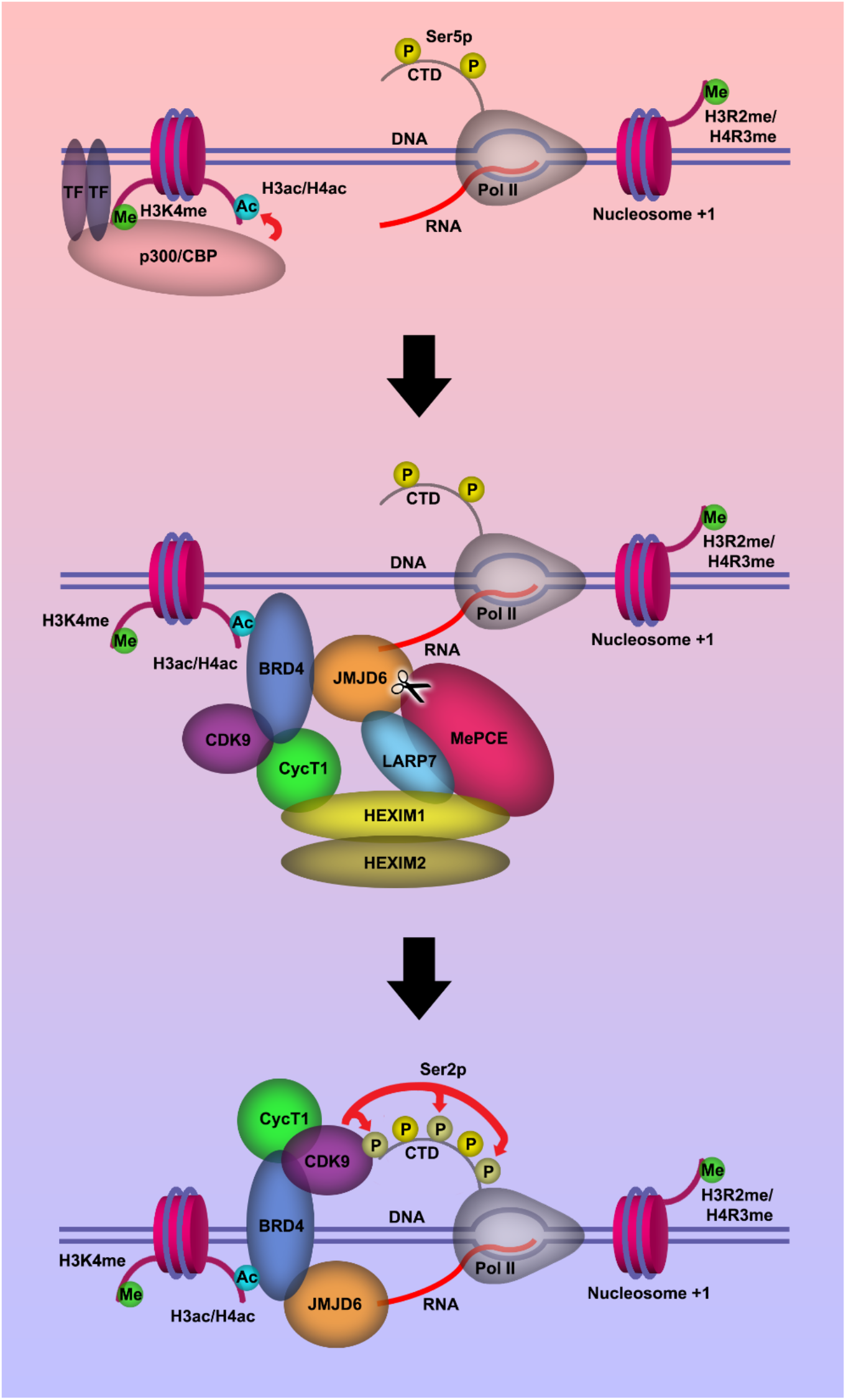
Model of P-TEFb release from 7SK snRNP complex. **a.** Pol II is initiated at the TSS and remains in the paused state until further stimulation. Transcription factors and H3K4me recruits p300/CBP, which acetylates and generates H3ac and/or H4ac. **b.** H3ac and/or H4ac recruits BRD4, which associates with 7SK snRNP/P-TEFb complex and JMJD6. JMJD6 associates with ssRNA from Pol II. JMJD6 digests MePCE to disrupt the 7SK snRNP complex to release P-TEFb (CDK9). **c.** BRD4/JMJD6 brings CDK9 in close proximity of CTD of Pol II. CDK9 phosphorylates Ser2 motifs on CTD of Pol II.

## Materials and Methods

### Protein expression and purification

The cDNA corresponding to gene of wild-type JMJD6 (1-343), inactive mutant JMJD6 (1-343) H187A/D189A/H273A/K204A/N287A, HEXIM2, and LARP7 was cloned into a pET28a vector containing an N-terminal His^6^ tag. The DNA corresponding to gene of JMJD6 (1-403) wild-type was cloned into a pGEX vector containing an N-terminal GST tag and TEV linker. All proteins were expressed in Rosetta (DE3) *Escherichia coli* cells. All cell cultures were grown to A600 value equal to about 1.0 and induced with a final concentration of 1.0 mM isopropyl 1-thio-β-D-galactopyranoside overnight at 16°C. Cells were resuspended in nickel-binding buffer (50mM Tris-HCl, pH8.0, 1M NaCl, 1mM PMSF) and lysed using a sonicator (Fisher Scientific Sonic Dismembrator Model 500) at 35% power, 10 sec ON, 5 sec OFF, for 20 min. The lysate was centrifuged at 16,000 rpm at 4°C for 30 min. The supernatant corresponding to His-JMJD6 (1-343) was loaded to 7mL of Ni-NTA resin (GE Healthcare), washed with nickel-binding buffer containing 20mM imidazole, and eluted with nickel-binding buffer containing 500mM imidazole. The supernatant corresponding to GST-tev-JMJD6 (1-403) was loaded to 7mL of glutathione agarose resin (Thermo Scientific), washed with nickel-binding buffer, and eluted with nickel-binding buffer containing 30mM glutathione. All eluted JMJD6 products were concentrated and purified on a superdex 200 10/300 GL column (GE Healthcare) previously equilibrated with nickel-binding buffer containing 15mM β-mercaptoethanol. The cDNA fragment encoding full-length MePCE and HEXIM1 with an N-terminal His-tag was cloned under control of the polyhedrin promoter into a previously described baculovirus transfer plasmid (Kozono et al., 1994). Recombinant virus was made by co-transfection into SF9 insect cells (Invitrogen) of the plasmid and BacVector3000 baculovirus DNA (Novagen) using the calcium phosphate co-precipitation method. High titer virus stock was prepared by infection of SF9 insect cells. MePCE and HEXIM1 protein was produced by infection of High Five insect cells (Invitrogen) at high multiplicity of infection. Four days later, the cells were lysed by sonication in nickel-binding buffer. Lysates were cleared by centrifugation (20,000 rpm, 60 min) and proteins were purified from the supernatant using 7mL of Ni-NTA resin (GE Healthcare). The protein was eluted from the column with nickel-binding buffer containing 500mM imidazole.

### Crystallization, Data Collection, structural determination, and refinement of JMJD6

His-JMJD6 (1-343) was crystallized by vapor diffusion in sitting drops with 0.1M sodium citrate pH 5.6, 1.0M ammonium phosphate monobasic at 8 ^o^C. The crystals were soaked in soaking buffer; 0.1M sodium citrate pH 5.6, 1.0M ammonium phosphate monobasic, 10mM mono-methyl arginine, 3.75mM αKG, 3.75mM Iron(II) sulfate. For data collection, His-JMJD6 (1-343) crystals were transferred to a cryo-protecting buffer (soaking buffer supplemented with 25% glycerol (v/v)) and frozen in liquid nitrogen. All data used in structure solving and refinement were collected on a beam line 4.2.2 (MBC-ALS) at the Advanced Light Source (Berkeley, ALS, USA). Data were integrated and scaled using the HKL2000 suite of programs. Structural determination and refinement results are shown in Table S1.

### Western blot analysis

To analyze protein levels, wild-type MEF, *Jmjd6* knockout (KO) MEF, wild-type JMJD6 overexpression in *Jmjd6* KO MEF, inactive mutant JMJD6overexpression in *Jmjd6* KO MEF, and MePCE overexpression in MEF cells were grown on 10 cm plates to be harvested and lysed using a standard RIPA buffer mixed with cOmplete Protease inhibitor cocktail (Roche). MEFs were generated from *Jmjd6^tm1.1Gbf^* knockout mice as previously described (PMID: 15345036, PMID: 21060799). Total cellular extracts in the presence of a protein standard (Bio-rad) were resolved by 8-12% gradient SDS-PAGE and transferred to a 0.22 μm nitrocellulose membrane and incubated with specific antibodies overnight at 4°C. Antibodies used in this investigation were: Anti-JMJD6 (Santa Cruz Biotechnology, sc-28348), Anti-LARP7 (Abcam, ab-134757), Anti-HEXIM1 (Santa Cruz Biotechnology, sc-390059), Anti-MePCE (Bethyl Laboratories Inc., A304-184A), Anti-Actin (Santa Cruz Biotechnology, sc-8432), Anti-His (Santa Cruz Biotechnology, sc-533073), Anti-Pol II CTD (Gift from Dr. David Bentley), Anti-Pol II Ser2p-CTD (Gift from Dr. David Bentley), and Anti-CDK9 (Santa Cruz Biotechnology sc-13130).

### *In vitro* MePCE cleavage assay

Full-length MePCE protein with N-terminal His-tag, mixed with EDTA-free cOmplete protease inhibitor (Roche), αKG, Zn^2+^, and HEPES pH 6.5, was titrated with recombinant wild-type JMJD6 and placed in 37 ^o^C for 2 hours. The reaction was subject to western blot analysis using monoclonal anti-His antibody (Santa Cruz Biotechnology, sc-533073). The reaction was reproduced in two separate experiments and the blot bands were quantified using ImageJ.

### Mass Spectrometry

MePCE (161-179) R171-me2s/C177S peptide, mixed with EDTA-free cOmplete protease inhibitor (Roche), αKG, Zn^2+^, and HEPES pH 6.5, was treated with recombinant wild-type JMJD6, inactive mutant JMJD6, or peptide alone and placed in 37 °C for 2 hours. 1 μL of reaction sample is mixed with 1 μL of a-cyano-4-hydroxycinnamic acid (10 mg/ml in 50% ACN, 0.1% TFA). The mixture is spotted on the MALDI target and allowed to air dry. The sample is analyzed by a Microflex-LRF mass spectrometer (Bruker Daltonics, Billerica, MA) in positive ion reflector mode. External calibration is done using a peptide calibration mixture (4 to 6 peptides) on a spot adjacent to the sample. The raw data is processed in the FlexAnalysis software (version 3.4.7, Bruker Daltonics) and exported in mzXML format. The mzXML files were analyzed on ProteoWizard. Data points were normalized to the intensity of the undigested peptide input and plotted on SigmaPlot v11.0. The reaction was reproduced in three separate experiments.

### Fluorescence Polarization experiment

Given the known structure of JMJD6, where aromatic residues are present in close proximity to the binding pocket (Y131, F133, W174, W272), the following tryptophan fluorescence assay was deemed suitable for characterizing various peptide binding activity. All the regular and methylated peptides were synthesized by AnaSpec Inc. (Histone 3) or Peptide 2.0 Inc. (MePCE). 5 μM JMJD6 (1-343) was titrated and equilibrated with fixed concentrations of each peptide respectively, incubated at 25°C for 30 min between each titration intervals, and subject to fluorescence measurement. The buffer used in the fluorescence quenching assay was 100 mM NaCl, 20 mM Tris-HCl pH 6.5, and 0.05% Tween-20. The excitation wavelength of 280 nm and the emission wavelength of 342 nm was used for data collection and recorded with a Fluoromax-3 spectrometer. The titration samples were prepared and analyzed in parallel as duplicates, triplicates, or quadruplicates. All values at different titration points were compiled, normalized against the maximum value obtained prior to titration and averaged. The error bars indicate the normalized minimum and maximum values at any given titration point. The K_D_ for each peptide was calculated by fitting to a 4 parameter sigmoidal dose-response curve with SigmaPlot v11.0.

### Microscale Thermophoresis (MST) experiment

His-JMJD6 (1-343) labeled with fluorescent NT-647 dye at a constant concentration of 100 nM was mixed with sixteen serial dilutions (~1.5nM–300μM) of peptides derived from MePCE (154-187), MePCE (154-184) R170-me2s, MePCE (154-184) R171-me2s, Histone 3 (1-21), Histone 3 (1-16) R2me2s, C-peptide (57-87). C-Peptide (57-87) was used as a negative control. MST experiment was performed using Monolith NT.115 (NanoTemper Technologies). His-JMJD6 (1-343) was titrated with peptides in PBS-T buffer (137 mM NaCl, 2.7 mM KCl, 10mM Na2HPO4, 1.8mM KH2PO4, and 0.05% Tween-20). The change in the fluorescence of bound and unbound labeled His-JMJD6 (1-343), ΔF, is indicative of the peptide binding. Plotting ΔF vs. peptide concentration facilitated the generation the dissociation curves, computed by the NTP program. The K_D_, reflecting the affinity of each of the peptides for His-JMJD6 (1-343), was obtained. The error bars indicate the normalized minimum and maximum values at any given titration point. Each experiment was performed in triplicate or quadruplicate.

### Modeling of JMJD6(1-343)-MePCE(164-178) interaction

Crystal structure of JMJD6 (1-343) monomer (PDB:6MEV) sans methylarginine was used as the template to build in residues corresponding to MePCE (164-178) near the catalytic center of JMJD6 in PyMol. No regard for clashes, bonds, or optimization was considered. Structure was exported in .pdb format and uploaded to YASARA energy minimization server (http://www.yasara.org/minimizationserver.htm) using default parameters (Krieger, Joo et al. 2009). Energy-minimized output model was converted to .pdb format and the MePCE (164-178) residues were minimally adjusted in PyMol to superimpose R171 of MePCE with the methylarginine observed in the crystal structure (PDB:6MEV).

### RNA-Seq

RNAs from wild type MEF cells, MEF cells with *Jmjd6* knockout, and MEF cells with JMJD6 overexpression cells in *Jmjd6* knockout background, respectively, were extracted with Trizol reagent (ThermoFisher Scientific). The extracted RNAs were then sent to Quick biology (Quick Biology, Pasadena) for further mRNA purification using oligo-d(T) beads. The purified mRNA was then used to build a mRNA library. Mouse Genome mm10 was used as the reference.

## Acknowledgement

We thank Dr. David Price from University of Iowa for cDNAs of MePCE and HEXIM1/2, samples of 7SK snRNP complex from Dr. Qiang Zhou at UC Berkeley, Dr. Peter Henson, Dr. James Hagman, Dr. Shaodong Dai, and Dr. Yang Wang, and Janice White for long time support in this project, and other researchers at National Jewish Health (NJH) for their kind support. Binding data was obtained from the Biophysics Core facility at Anschutz Medical Center, University of Colorado at Denver. Mass spectrometry data was obtained from the Proteomics and Metabolomics facility at Colorado State University. S.L. is supported by NIH Training grant T32AI007405-28 (to P. M.). H.L. is partially supported by NIH Training grant 5T32AI074491-07(to J. C.). G.Z. were partially supported by CA201230 (to K. B.), and NJH bridge fund.

---

**Figure S1.**
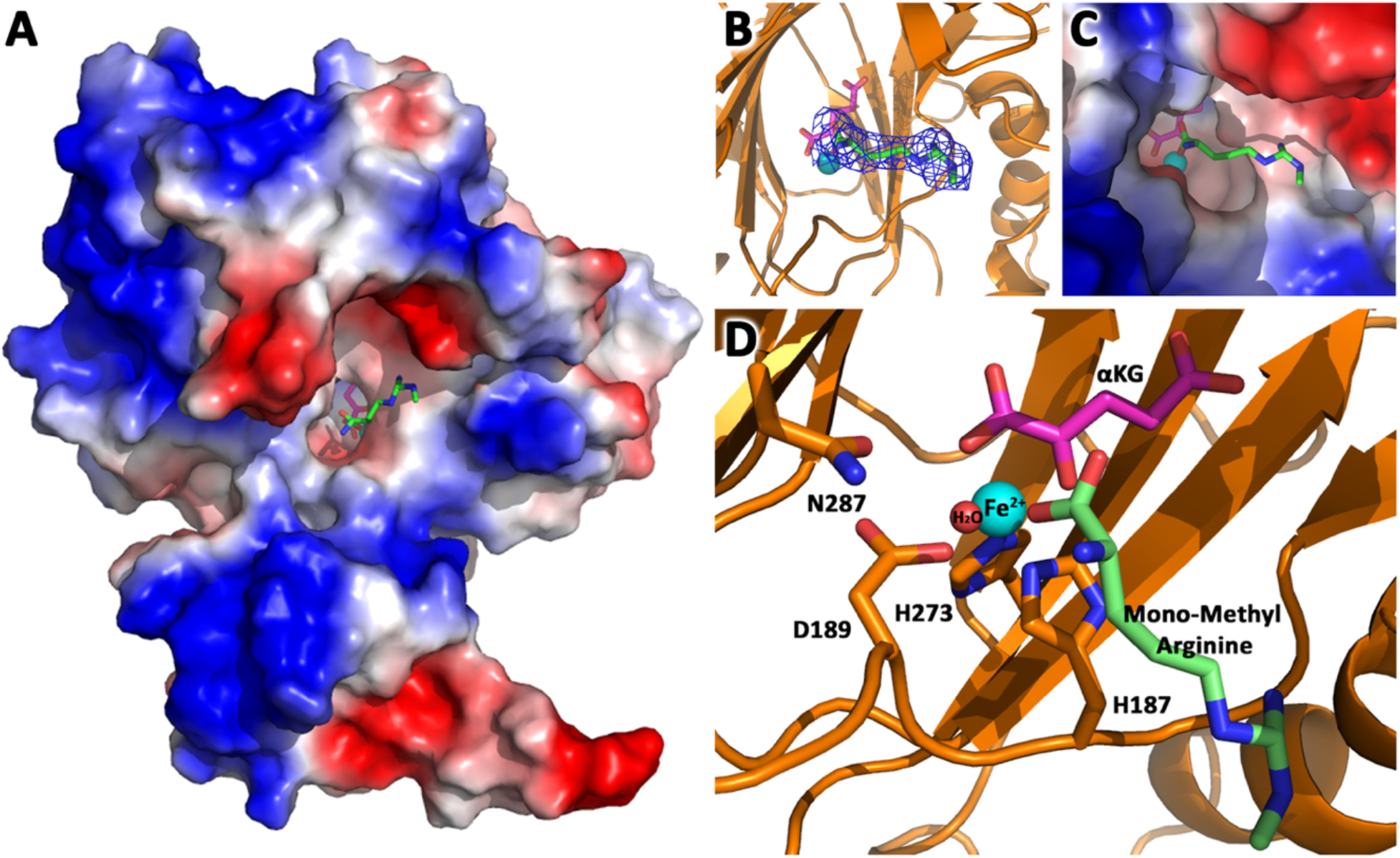
JMJD6 binds to monomethyl arginine (2-of-4). **a.** complex structure of JMJD6 (1-343) and monomethyl arginine (MM-Arg). Surface charges were generated using PyMOL (Action > generate > vacuum electrostatics >protein contact potential) (https://pymol.org/2/). Red represents negatively-charged surface, Gray represents neutral-charged surface, and Blue represents positively-charged surface. **b.** Omit map 2Fo-Fc electron density of MM-Arg. **c.** Magnified view of MM-Arg in the catalytic center of JMJD6 **d.** Coordination of elements at catalytic center.

**Figure S2.**
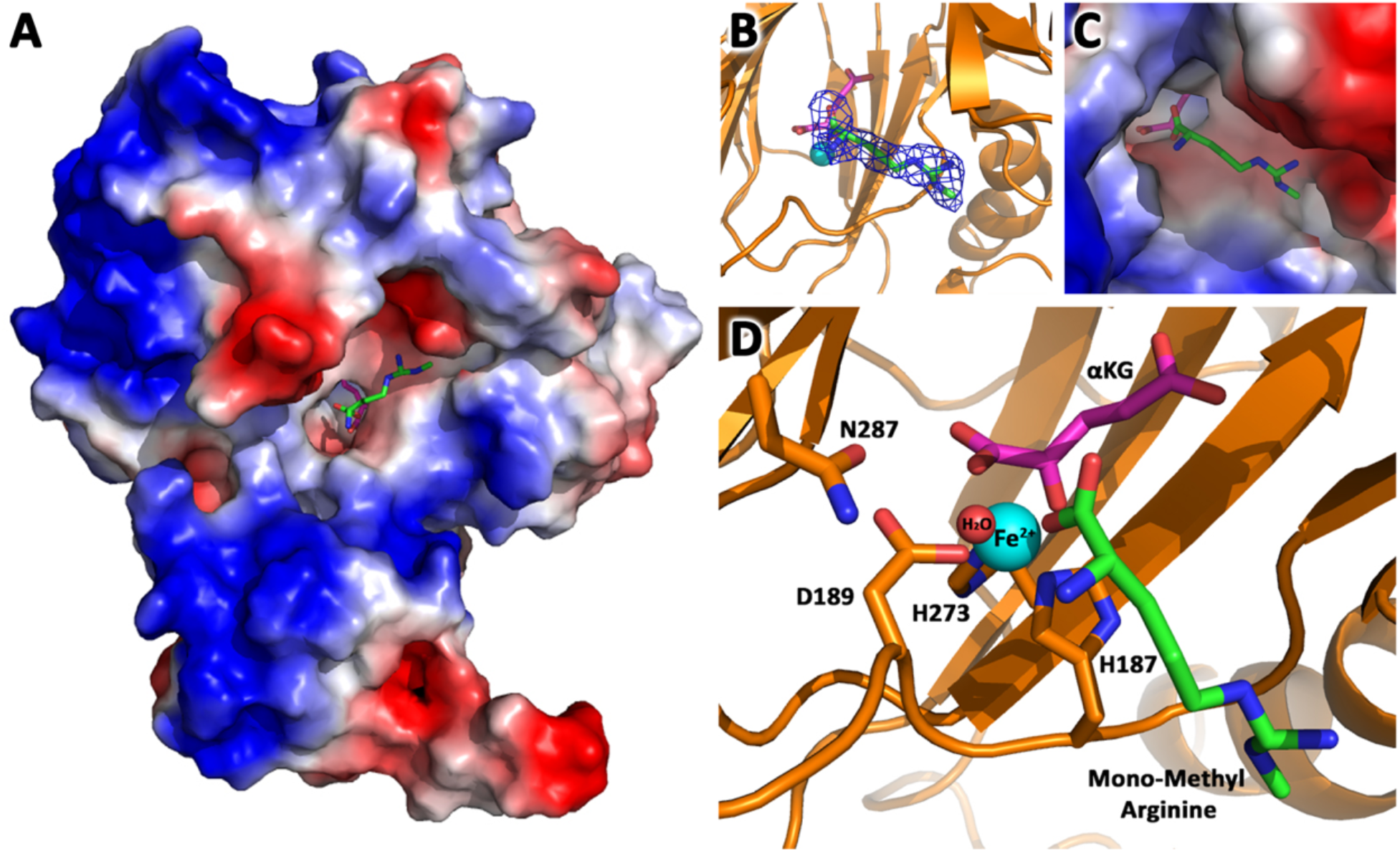
JMJD6 binds to monomethyl arginine (3-of-4). **a.** complex structure of JMJD6 (1-343) and monomethyl arginine (MM-Arg). Surface charges were generated using PyMOL (Action > generate > vacuum electrostatics >protein contact potential) (https://pymol.org/2/). Red represents negatively-charged surface, Gray represents neutral-charged surface, and Blue represents positively-charged surface. **b.** Omit map 2Fo-Fc electron density of MM-Arg. **c.** Magnified view of MM-Arg in the catalytic center of JMJD6 **d.** Coordination of elements at catalytic center.

**Figure S3.**
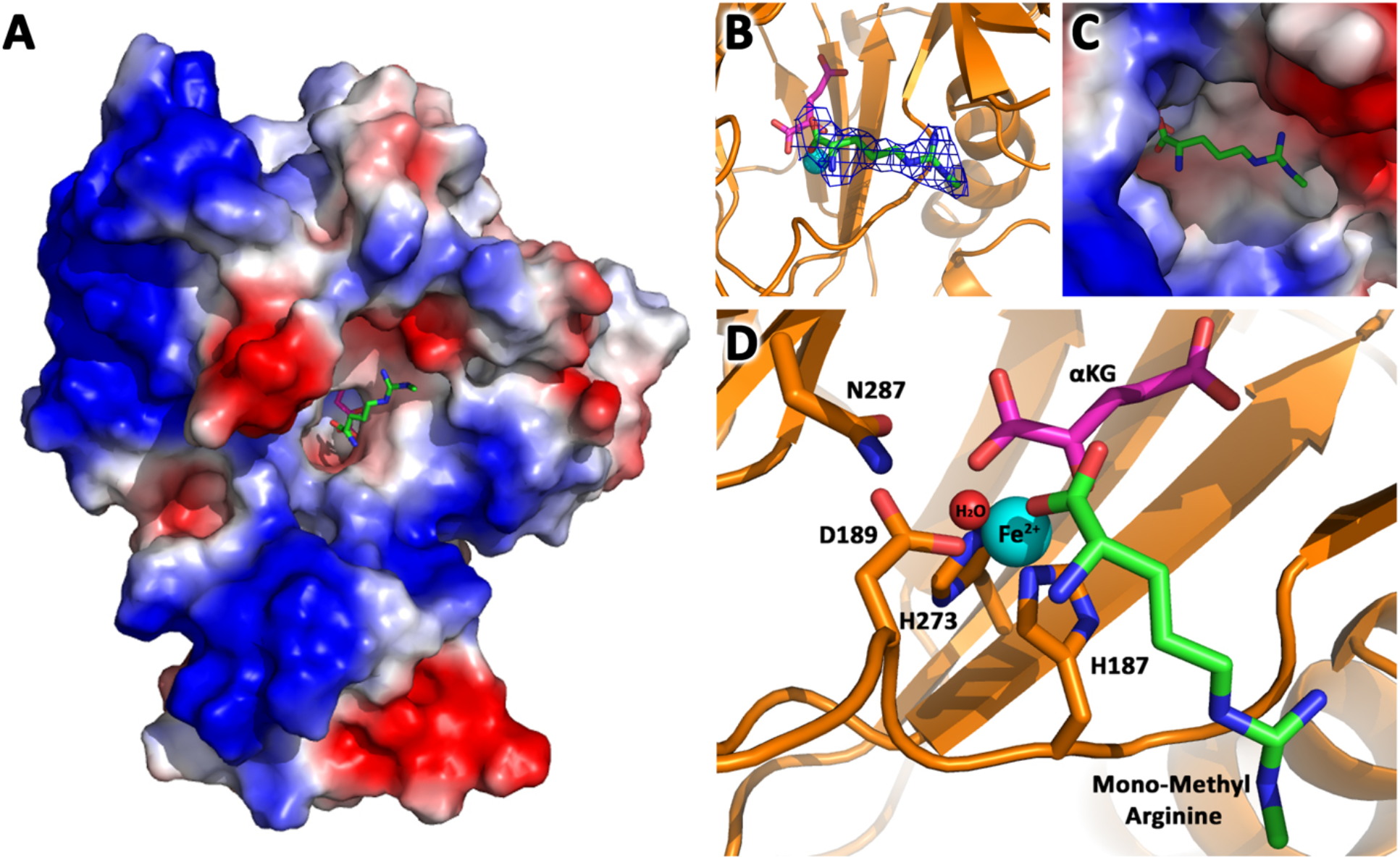
JMJD6 binds to monomethyl arginine (4-of-4). **a.** complex structure of JMJD6 (1-343) and monomethyl arginine (MM-Arg). Surface charges were generated using PyMOL (Action > generate > vacuum electrostatics >protein contact potential) (https://pymol.org/2/). Red represents negatively-charged surface, Gray represents neutral-charged surface, and Blue represents positively-charged surface. **b.** Omit map 2Fo-Fc electron density of MM-Arg. **c.** Magnified view of MM-Arg in the catalytic center of JMJD6 **d.** Coordination of elements at catalytic center.

**Figure S4.**
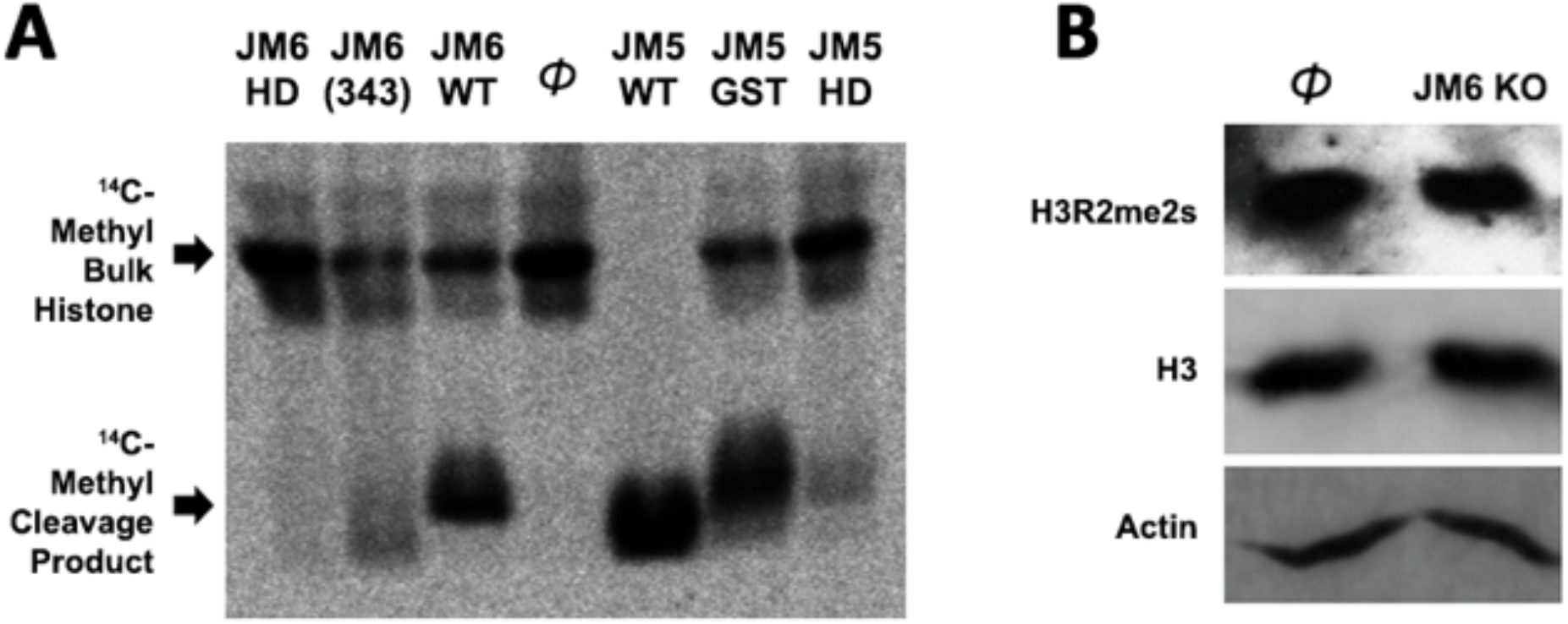
Histone tails are not the cognate substrate of JMJD6. **a.** ^14^C-labeled bulk histone reaction and resultant cleavage product when treated with JMJD6 and JMJD5 *in vitro*. **b.** JMJD6 knockout has no effect in the content of both arginine methylated H3 and total H3 in MEF cells *in vivo*.

**Figure S5.**
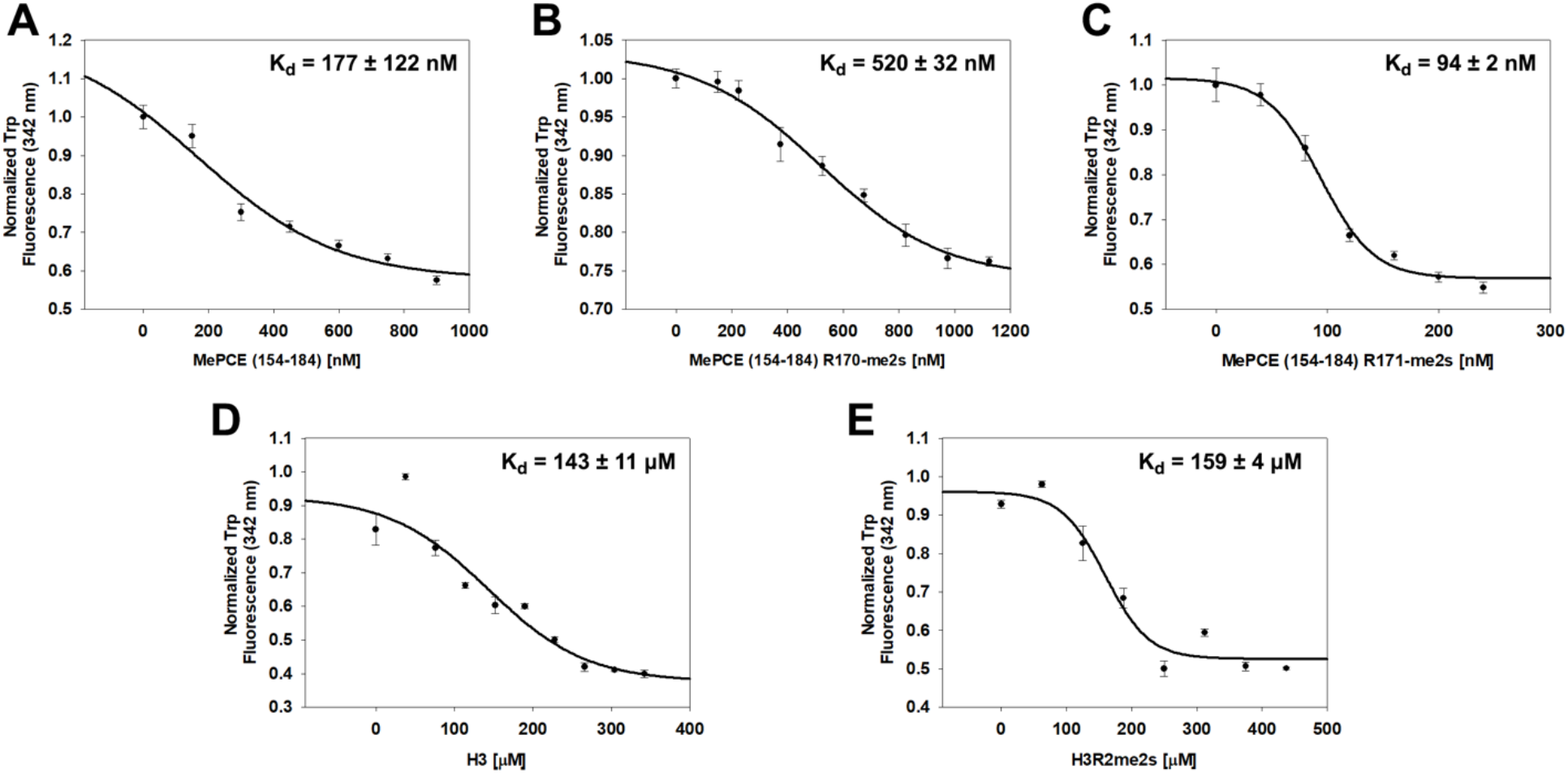
JMJD6 specifically binds to MePCE site containing residues 164-178 (determined via fluorescence polarization). **a**. The binding of His-JMJD6 (1-343) to unmodified MePCE (154-184). **b**. The binding of His-JMJD6 (1-343) to MePCE (154-184) R170-me2s. **c**. The binding of His-JMJD6 (1-343) to MePCE (154-184) R171-me2s. **d**. The binding of His-JMJD6 (1-343) to unmodified Histone 3 (1-21). **e**. The binding of His-JMJD6 (1-343) to Histone 3 (1-16) R2-me2s.

**Figure S6.**
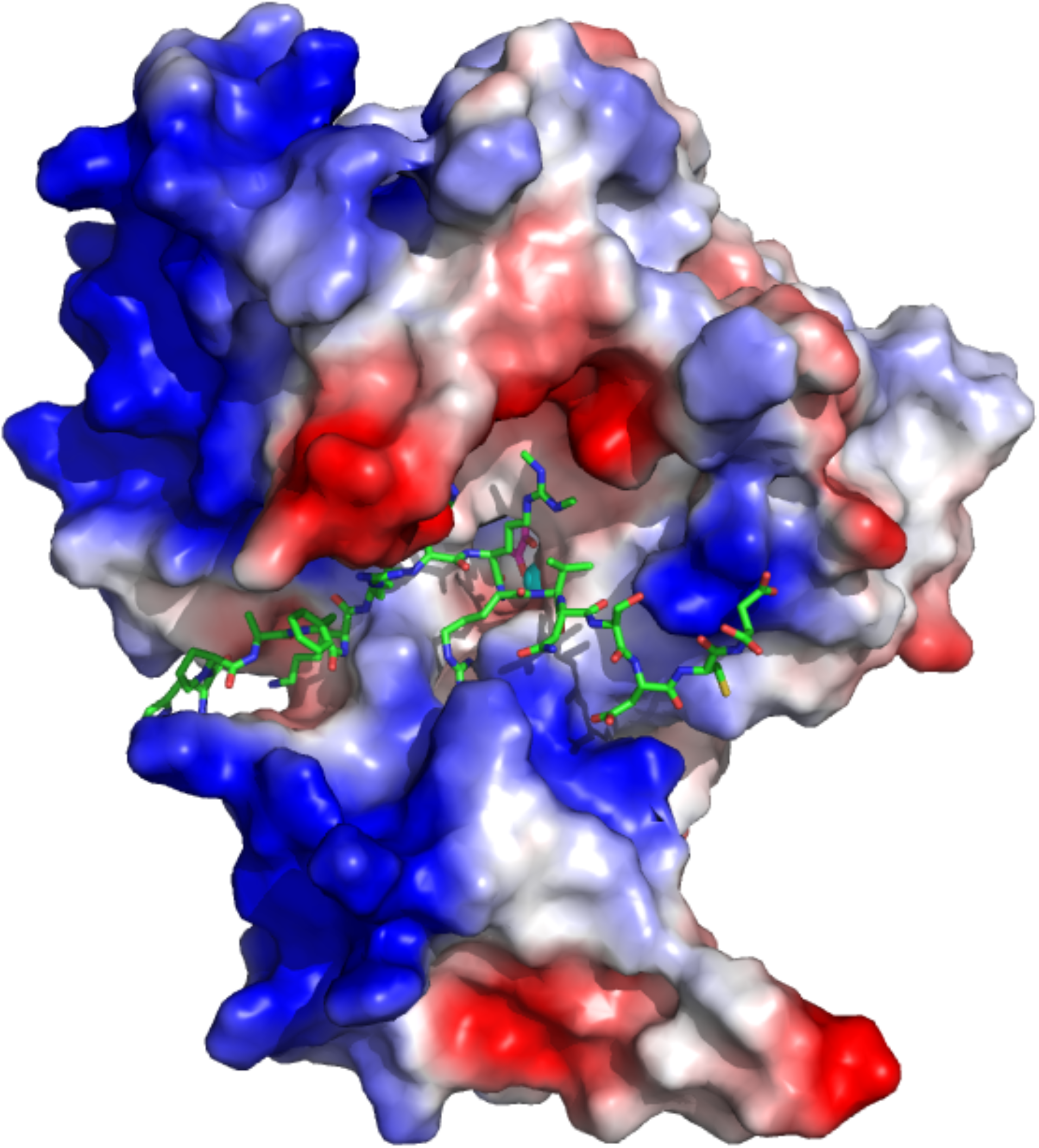
Computational complex structure model of JMJD6 (1-343) and MePCE (164-178) derived from YASARA Energy Minimization server. Minor spatial adjustments were made on R171me2s of computational model to better overlap with experimental model.

**Table S1.**
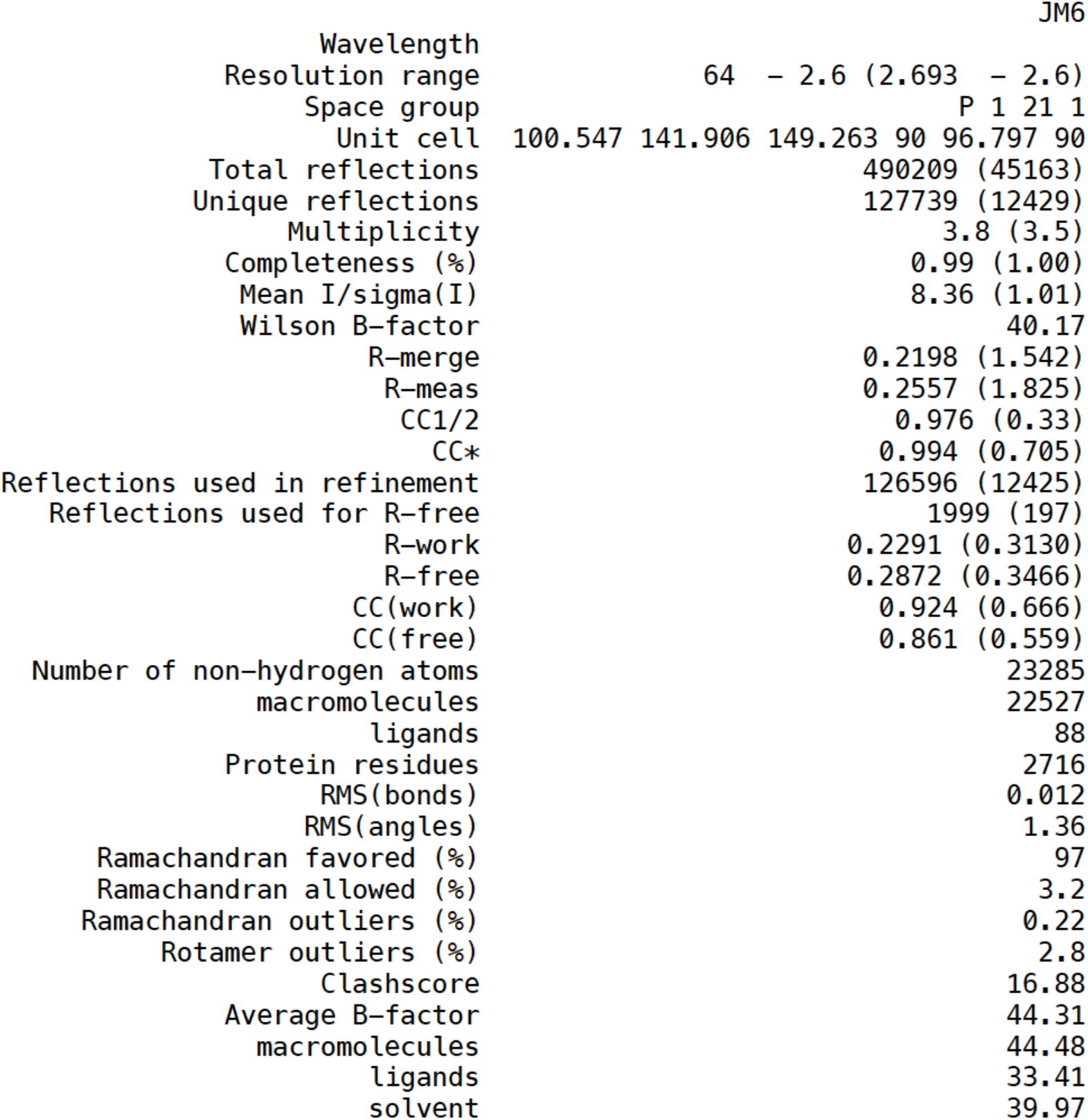
Crystallographic statistcs of complex structure of JMJD6 and a methylated arginine.

**Table S2. Protein Composition Analysis via Mass Spectrometry**. Bacteria expressed and purified JMJD6 are subjected to mass spectrum analysis, all potential contaminated trace protein candidates are listed. No known protease candidate is identified from the list.

**Table S3. Total RNA-seq reads of wild-type MEF, JMJD6 knockout MEF, and JMJD6 overexpressed in JMJD6 KO background MEF**.

